# Representation and Retrieval of Brain Connectivity Information derived from TMS Experiments

**DOI:** 10.1101/2023.01.22.522249

**Authors:** George F. Wittenberg, Xiaoqi Fang, Souvik Roy, Bryan Lee, Nataša Miškov-Živanov, Harry Hochheiser, Layla Banihashemi, Michael Vesia, Joseph Ramsey

## Abstract

**Background:** Transcranial magnetic stimulation (TMS) is a painless non-invasive method that allows focal activation or deactivation of a human brain region in order to assess effects on other brain regions. As such, it has a unique role in elucidating brain connectivity during behavior and at rest. Information regarding brain connectivity derived from TMS experiments has been published in hundreds of papers but is not accessible in aggregate.

**Objective:** Our objective was to identify, extract, and represent TMS-connectivity data in a graph database. This approach uses nodes connected by edges to capture the directed nature of interregional communication in the brain while also being flexible enough to contain other information about the connections, such as the source of information and details about the experiments that produced them.

**Methods:** Data related to interregional brain connectivity is first extracted from full-text publications, with creation of a table-like structure that list data of multiple types, principally the source and target brain regions, sign (excitatory/inhibitory) and latency. While machine-reading methods were explored, so far human experts have had to extract and verify data. These data are used to populate a `neo4j` graph database. A graphical user interface coupled with a query system allows users to search for networks and display information about connections between any two brain regions of interest.

**Results:** Experiments involving two TMS stimulating coils, in which one is over a putative source region and the other is over another region with a measurable effect in the body (such as the primary motor cortex) are the most straightforward to represent in the database. Even in those experiments, differing conventions for naming regions, and differing experimental parameters such as stimulation intensity and coil position, create issues for representing data from multiple studies in the same database. Other types of experiments, such a neuromodulatory stimulation such as rTMS, can provide information regarding connectivity, but are harder to represent. But we have, thus far, stored information about 100 connections in the database and demonstrated its utility in exploring direct and indirect connections between brain regions. We have also explored adding a causal inference method to those connections, allowing information about latency to shape the connections retrieved given latency limits.

**Conclusion:** A graph database can flexibly store information about interregional brain connectivity and is particularly useful for exploring the temporal aspects of brain networks.

## 1. Introduction

### 1.1. Transcranial magnetic stimulation contribution to brain studies

#### 1.1.1. TMS as probe of function

Transcranial magnetic stimulation (TMS) is a method based on Faraday ‘s Law of Induction and involves non-invasive induction of brief electrical currents in the brain that can be sufficient to cause action potentials in local neuronal elements, or otherwise affect membrane potential. TMS devices consist of a power supply that charges a capacitor, whose stored energy is discharged through a coil. This method became practical with the development of high-voltage silicon-based switches to control the flow of current. Some of the earliest studies with such modern devices used TMS to study the motor cortex, as single pulses of TMS over the primary motor cortex (M1) can cause easily measurable motor-evoked potentials (MEP) in skeletal muscles. The MEP amplitude represents an excitability measure of a large population of neurons. In a sense, these are some of the simplest ways in which TMS can be used as a physiological read-out to measure *connectivity*. The connection of the upper motor neurons, in the brain, to the lower motor neurons, in the spinal cord, and from them to the muscles is measured, with determination of several parameters: threshold, latency, magnitude of response, and specificity (which muscles are activated.) Intracortical and spinal processes can be measured through paired-pulse stimulation, using the MEP as the outcome measure, with an assumption that the first pulse conditions the response of the brain or spinal cord to the second. Such measures in the understanding of human brain physiology and pathophysiology have been reviewed [1, 2, 3, 4, 5].

#### 1.1.2. TMS as probe of connectivity

The place of TMS experiments in the understanding of brain connectivity has been reviewed [6]. With two TMS coils, one over M1 and one over another brain region, the effects of the other brain region on M1 can be measured, reflecting the connectivity of the two brain regions. In a way analogous to MEP measurement, information about threshold, timing, and magnitude of modulation can be collected. In these experiments, different values of interstimulus interval and stimulation strength can be used, and stimulation strength is often normalized to the motor threshold for response in a hand muscle to stimulation of M1. So data about connectivity is about responses over a range of parameter space that contains the most effective interstimulus interval and strength of stimulation for modulation of the MEP. While in some cases the effect of the region tested on M1 will be through a direct connection, in an intact brain the effects could be mediated by any number of other regions, limited only by the conduction delays expected between different regions. *Triple coil* experiments can also be done, testing the modulatory effect of one region on the modulation of M1 by another region.

A great strength of TMS lies in the temporal precision with which stimulation is delivered. So for two and three coil experiments, the relative timing of the stimuli is precise and the effect of changing the relative timing can be assessed. One limitation is the time it takes to explore the variety of timings, as stimulators and the nervous system need time to recover, requiring a wait of several seconds between trials.

Other types of experiments that give information about connectivity include TMS combined with a functional imaging modality such as fMRI, PET, EEG, or fNIRS. In these type of experiments, the local and remote effects of stimulation are measured through modalities sensitive to change of activity. This can be done at rest, but also during behaviors, as allowed by the imaging modality. Each modality has different limitations in terms of sensitivity to TMS-induced artifacts, temporal resolution, spatial resolution, and access to deep brain structures. But such studies can provide a fuller picture of connectivity of the target region because they all provide information regarding responses in multiple potential connected regions.

Finally, there are modulatory approaches to acquiring information regarding connectivity. With these approaches, repetitive TMS (rTMS) or transcranial direct current stimulation are used prior to other types of connectivity analysis. The lasting (on order of minutes) effects of target neuromodulation on connection values can be assessed. This can also be done in reverse fashion, with connectivity assessments used to predict response to neuromodulation [7].

### 1.2. Constraints on TMS Measures of Interregional Brain Connectivity

The degree to which brain regions are connected is constrained by anatomical factors, essentially the white matter *skeleton* of the brain. While this skeleton has been defined, it is probabilistic, as tractography relies on probabilistic methods to trace pathways. Ideally, known anatomical connectivity would constrain the connectivity derived from TMS experiments. Practically, this is complicated by uncertainly of the existence of a direct pathway in any given individual. But other information, such as latency of stimulation effects, provide clues as to the directness of interregional connections.

#### 1.2.1. Implying Causality in a Network

##### Network issues

With a two-coil experiment, the goal is measuring the effect of stimulation one area on the function of a second. The implied model is a direct connection between the two areas. However, both regions are parts of multiple brain networks with ongoing activity, and stimulation occurs in the context of those connections and connectivity. This complexity can be addressed by measuring only shorter latency effects, control of behavioral state, and, potentially through consideration of other known connections. Sophisticated approaches to modeling causality in complete networks have been developed [8] and applied to connectivity derived from fMRI [9].

#### 1.2.2. Machine Readers

Machine readers are tools that allow scalable interaction inference from literature for building network models. They identify entities and events, as well as their relationships [10, 11]. The state-of-the-art automated reading engines are capable of finding hundreds of thousands of events and relationships from thousands of papers, in a few hours. Both entities and events can be represented as graph nodes, whereas their relationships are represented as graph edges [12]. For this work, we selected to explore the TRIPS reading engine, which is a semantic parser with broad domain coverage. TRIPS was developed with the goal to be domain general, and it produces logical forms grounded in a general ontology [13, 14]. In contrast to many other semantic parsers, which are limited to simple domains and not transferable to new domains, the TRIPS parser has been demonstrated to work well in diverse domains, incorporating domain-specific named entity recognition where needed.

#### 1.2.3. Publication limitations

Whether humans or automated agents read a paper describing the use of TMS to study connectivity, there are many limitations on the efficient, reliable, and complete extraction of data from that paper. Publications have usually not followed any guidelines for uniform representation of experiments – in fact none exist in the TMS sphere, to our knowledge. This is an issue that we hope to address by developing recommendations and proposing structured methods for providing access to raw data. Chief among these limitations is the varied terminology for describing brain regions, experiment parameters, and effects. This has caused problems for machine readers in a way more significant than for other types of dynamical biological networks, for which machine readers have been used to extract information [15]. The most relevant data is often present in graphs, so one approach for extraction would be of reverse plotting applications [16]. And while numbers of individuals, means, and standard deviations are often reported, generally each individual ‘s data set is not extractable from a paper. Other issues include variation in behavioral paradigms, TMS simulator strengths, referencing of stimulation strength to different standards, such as active or resting motor thresholds, and multiple methods for control, including use of sham coils, or control stimulation locations.

### 1.3. Graph Databases

The network of interconnected brain regions maps naturally onto a graph database, the use of which has been increasing in bioinformatics [17]. Related to graph theory, the graph database is structured as a collection of *nodes* and *edges*. Information can be stored at both nodes and edges. The graph concepts map naturally onto brain circuits at all levels of analysis. For instance, neuron cell bodies can be considered nodes and neurites edges. In the macro scale that we are concerned with, they can be grey matter regions and white matter tracts. Graph theoretical representations of brain regional connectivity has been used to determine connectivity patterns derived from EEG and fMRI, leading to the *small world* concept of optimized white matter length and processing time [18].

## 2. Methods

We approached the problem of extraction of connectivity data and representation in a database first conceptually, by first drawing graphical representations of the connectivity implied in selected papers. We developed and improved an extraction method for individual data points. Each data point was a list of variables that included references to the paper, source and target nodes, timing information, and experimental descriptors. The columns of the database currently have the headings shown in table 1:

**Table 1.** Data Fields Extracted. These were the initial fields used to populate the database regarding individual connections, between source and target regions. Additional columns have been proposed, including ones that contain information about age, whether healthy or affected by a disease state, etc. example data from [19]

| Content | Example |
| --- | --- |
| Description of connection | rPMv-lM1 |
| Brain region – source of connection | rPMv |
| Target region stimulation strength | 110% RMT |
| Stimulation pattern | Single pulse |
| Coil orientation | posterior-anterior |
| Brain region receiving connection | lM1 |
| Source region stimulation strength | 100% RMT |
| Stimulation pattern | Single pulse |
| Coil orientation | posterior-lateral |
| Sign of effect (inhibition, facilitation or null) | Facilitation |
| Latency | 8 ms |
| How effect was measured (MEP, MRI, etc.) | MEP |
| Study type (Dual coil, neuromodulation, etc.) | Paired Coil TMS |
| Behavioral context (if any) | Target Presentation |
| Number of participants | 14 |
| Initials of person entering data | SR |
| Unique publication identifier | 10.1523/JNEUROSCI.4882-09.2010 |
| Journal, vol., pages | J Neurosci. 2010 Jan 27; 30(4) 1395-1401 |
| First Author | Buch |
| Publication year | 2010 |
| Free text regarding connection | measured in multiple behavioral states |

**Table 2.**
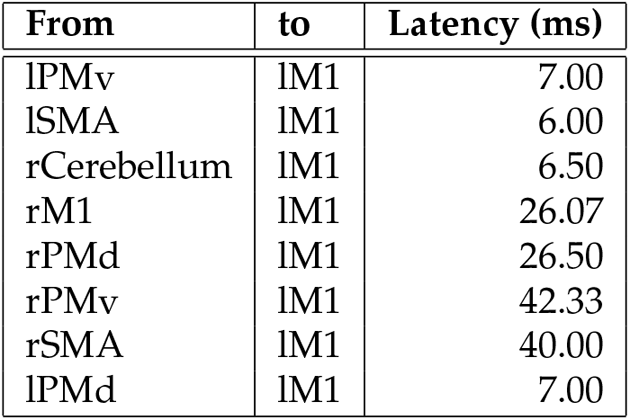
Eight connections to left M1 specified in dataset. The latencies represent averages when there was data from multiple papers.

**Table 3.**
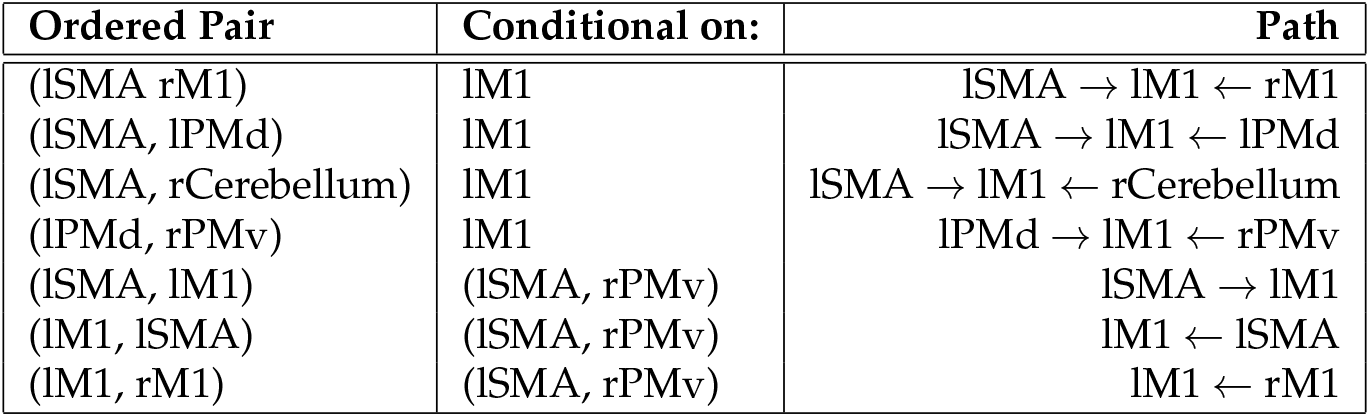
Example T-connection related to left M1. The latencies represent averages when there was data from multiple papers.

### 2.1. Literature Search & Machine Reading

Literature search was performed using PubMed[20], with search terms described in the Results section. Fulltext papers were downloaded and collected. As mentioned above, we selected the TRIPS-DRUM parser[13, 10] to process the data. TRIPS (<u>t</u>he <u>R</u>ochester <u>I</u>nteractive <u>P</u>lanner <u>S</u>ystem – originally used to provide transportation information) uses knowledge of language that includes statistical properties and grammar to extract *events* that correspond to relationships among entities in a tree-like logical form, similar to sentence diagrams used to teach grammar. TRIPS has been applied in the DRUM (Deep Reader for Understanding Mechanisms) project to “read papers and combine research results of individual studies into a comrehensive explanatory model of a complex mechanism” [10]. Using TRIPS-DRUM, we were able to identify potential data of interest, but not to extract all the data required to populate the database.

### 2.2. Human Extraction of Published Data

Because the machine readings approach has so far not led to extraction of all of the required fields, individuals, including medical and neuroscience students, extracted data regarding each connection studied in a paper, with review by the database coder and senior investigators for each piece of data. Once the methodology of the experiment was correctly identified, multiple lines of data could be extracted for that experiment. In some papers, there were multiple experiments, which required care to correctly match each experiment type with extracted data. Each unique combination of stimulation parameters and other conditions, such as experiment type, resulted in a single line in this *flat* database.

#### 2.2.1. Naming Convention

Naming brain regions is complicated by the many naming conventions used in human neuroscience. We used the NeuroNames [21] system, making a table of common synonyms in human motor neuroscience, to translate regional names not in that ontology. The graph database structure was designed to handle hierarchies of brain regions, for example the difference between body part representations within M1, and the multiple regions in the intraparietal sulcus. The goal was to name regions by side (left or right) and by the acronym or name in the NeuroNames list, such as PMD for dorsal premotor area, which has the unique identifier ‘3164.’ This was accomplished by finding the region name in the paper, and searching in NeuroNames.

### 2.3. Graph Database

We developed a web-based user interactive database application that allows information exchange between client and server (Figure 2.) This application was designed as a multi-layer system consisting of a REST API layer for Web service interfaces and a database layer for data storage. The server side was constructed using *flask*, a python micro-framework, since it is simple and extensible, embedded with only the core components for web development such as routing, request-handling and unit testing, while providing compatibility with numerous extensions, ensuring its feasibility for future scaling up. Several components were combined at the server end for the proper functioning of this application. The requests submitted by the client are first transmitted and handled by the REST API layer based on HTTP standard. The implementation details are hidden from the clients for simplicity and information security. We incorporated an in-memory cache layer with *Memcached*, a distributed memory object caching system, to reduce the number of requests handled by the primary database[22]. The application also utilized Apache *Lucene* to construct a full-text index system for tokenization, improve the accuracy and efficiency for indexing and searching[23]. An embedded data management system tracks the performance of components mentioned above to ensure the reliability and overall healthiness of the application. We also created an interactive visualization panel with *Popoto*, a d3-based graph visualization library, to represent the data retrieved from the underlying Neo4j database and provide intuitive representations of the brain circuit data at various levels.

### 2.4. Retrieval & Display

Users can simply click the nodes or enter textual queries to access the data, and explore the circuit knowledge database through the graphical displays obtained(Fig. 1). We will collect information on query statistics, including number, region names entered, and edge data accessed. We will be able to display the TMS-derived connections of key motor areas, such as the SMA (supplementary motor area), along with information on number of studies, consistency/conflict, and timing, of each potential connection to another area. Differences between networks from the normal state as a results of neurological disease and aging will be presented. Visualization is through multiple methods, including displaying networks on a brain template such a distorted version of the brain flat map from [24] with graph nodes close to the flat map representation of the source area to which it refers. Basic ball-and-stick networks such as we have developed already and 3-D visualizations in standard atlas space will also be available.

### 2.5. T-connectivity

Parallel to the work described above, data from the flat database originating from two-coil experiments could be used to perform causal modeling as has been done previously with graphs [25], but with an added *time constraint*. Data (either from the database itself, or synthetic data created to test the model) were given as input to a new algorithm in the TETRAD software system to derive what we are calling *t-connected* relationships implied by the conditional independence facts garnered from the experiments themselves. Since path latency information is available in the data, with the sign of the effect for each stage, and may be summed along paths, total latencies along paths are tracked, with the overall sign of the paths recorded as well, so that the excitatory connections would not change the sign of the expected effect, but inhibitory ones would. The system provides a summary of all potential paths consistent with this data, from one region to another, along with expected times and signs of effects along each possible path. In this way, it is possible to determine which paths can be followed in the experiment within a certain time limit, observed in the experiment. Thus, possible paths explaining a connection, across TMS experiments, are able to be extrapolated. This is not meant to predict exactly which paths in the experiment are in fact responsible for an effect given an stimulus, but rather to elucidate which paths *could* possibly be responsible for such an effect, given what is known currently in the database of experimental TMS results. As more experiments are added to the database, such query results may be refined, making this a potentially useful tool, summarizing existing experimental knowledge, but allowing time constraints to limit results to those that can explain time-limited behavioral observations.

## 3. Results

### 3.1. Literature Search & Machine Reading

#### 3.1.1. Literature Search

PubMed literature searches were carried out with the search terms (transcranial magnetic stimulation) AND (connectivity) resulted in 1936 retrievals on 3 Jan./ 2022. Requiring “connectivity” to be a title word reduced the number to 424, and further requiring “stimulation” to be a title word reduces it to 191, the papers screened out have about 6% papers that could provide data regarding connectivity. Some relevant papers use “TMS” in the title, or otherwise fail to use the term “stimulation” even if that is the basis of the connectivity measurement. Papers were selected individually from this list manually if the abstract mentioned a measure of connectivity being measured.

#### 3.1.2. Machine Reading

One test of machine reading came about with application of the TRIPS DRUM reader to three papers that were also manually processed [26, 27, 28]. The output of the reader correctly identified “events” with descriptions such as: Activate, Bind-interact, Decrease, Increase, Interact, Modulate, Stimulate, and Transform. These events could helpful in identifying statements in the papers regarding connectivity. For instance, the modulation of MEP size with different cerebellar-M1 stimulus intervals was identified in the text phrase “increased and decreased MEP” with the event *affected MEP* automatically extracted. However, identification of the affected target was only made four out of 29 times, and *causal parent* event never identified. The most useful part of the machine reader output was, thus far, the identification of text phrases that helped indicate if the paper was of interest for manual extraction.

### 3.2. Human Extraction of Flat Database

In parallel with the machine reading experimentation, students extracted information specified in the columns of the flat database. At the time of writing, there were 101 entries in the flat database, from 45 papers. The entries represent a diversity of experiment types, two coil experiments, behavioral perturbation, fNIRS, etc. Guidance was required in the interpretation of parameters, particularly when it came to timing, but names of areas were always found in the NeuroNames database. There are different types of time measures in TMS experiments; the one we cared most about was *latency*, the time between stimulation of one area and response in another. For a two-coil experiment, this measure is equal to the interstimulus interval. But there are also repetition intervals for each stimulation, interpulse intervals for trains of stimulation, duration of rTMS effects, for example. There is particular difficulty in interpretation of the intervals between stimulation and *behavioral* effect, since these involve a time period longer than the interregional latency.

### 3.3. Storage in a Graph Database

The flat database was read to create a graph database, as described in the Methods. Each line of the flat database was used to specify a node, with edges linking not only the regions that were connected, but to other information about that line of data, including the sign of the connection effect and the provenance of the data (the DOI of the publication.) This allowed more flexibility in representing multiple versions of the same connection from different experiments, and kept the data about each connection tethered to its source.

### 3.4. Retrieval & Display

Retrieval of information about connections can be accomplished by either using a cursor to explore the graphical interface which allows edges to displayed by clicking on the visual edge of each brain region ‘s representation, or through a text-based Cypher query Fig. 1. (Clicks on the graphical representations are converted into Cypher queries, which are SQL-like in structure, and are internal to the neo4j system [29].)

**Figure 1.**
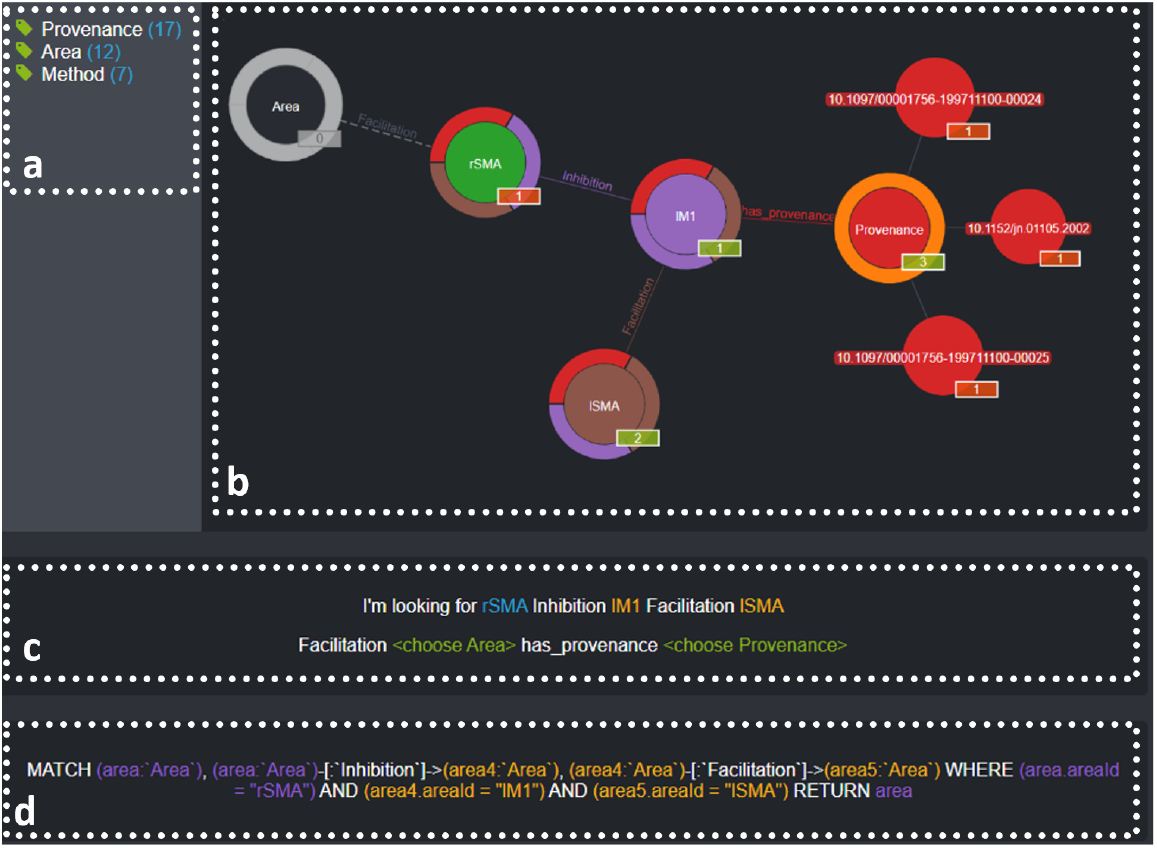
Query Page contains. **(a)** overview of type and number of nodes **(b)** graphical representation of queried data. Here rSMA (the green circle) is the source region for the query. The area lM1, inhibited by rSMA is *unfolded* by clicking at the purple section of ring around it. Similarly, lSMA is unfolded when clicking at the brown section of ring around lM1. The red section of ring stands for **Provenance**, and three publications represented as red circles with their DOIs provide evidence for inhibition effect from rSMA to lM1. **(c)** summary of the effect flow between brain areas **(d)** the Cypher (SQL-like) query automatically generated by clicking at the nodes and edges in (b)

**Figure 2.**
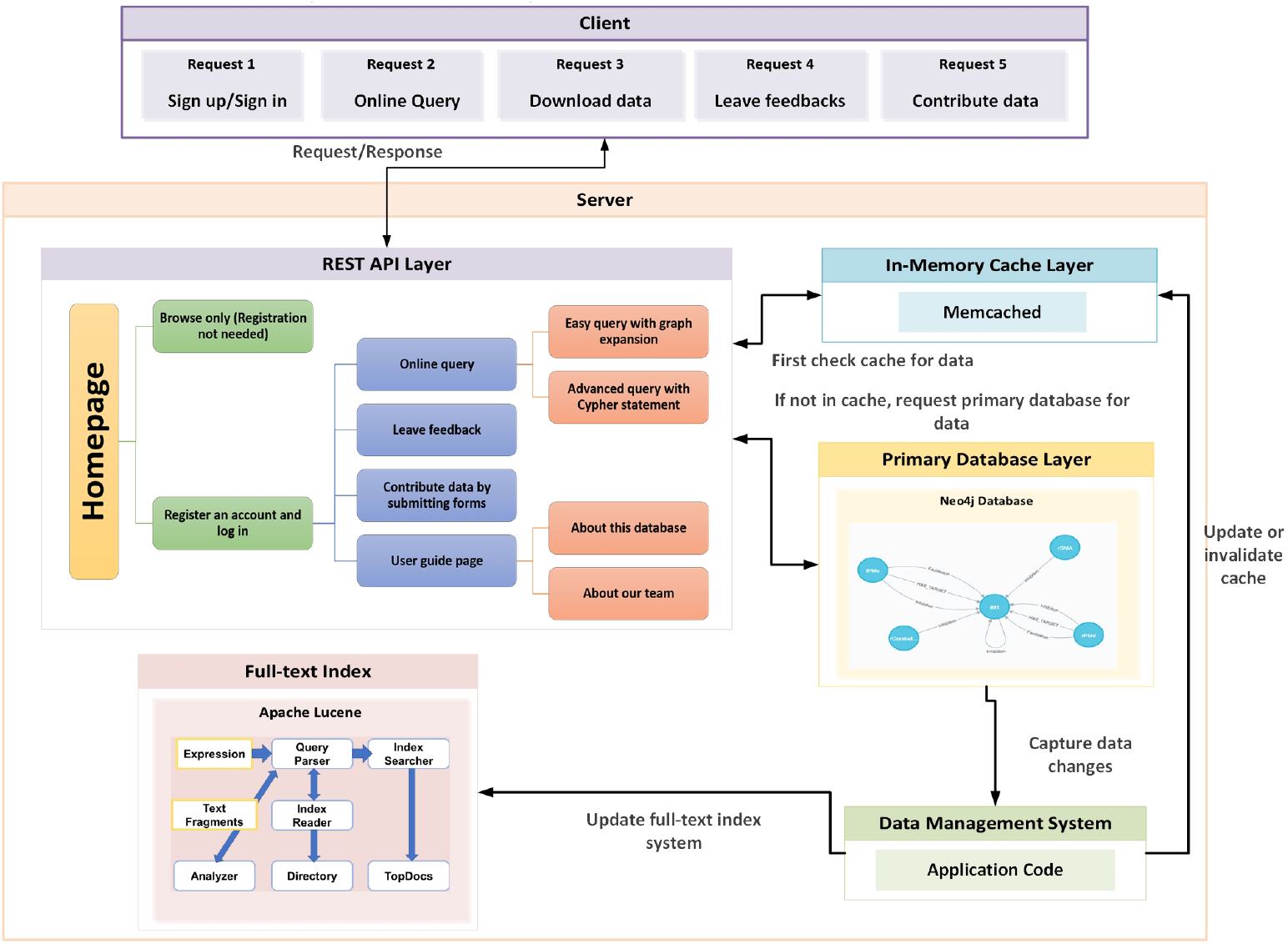
Application Architecture for showing the components of this application. Requests are submitted from the Client side, then received and processed by the REST API layer of Server side.

### 3.5. T-connectivity

As described in the Methods section, selected data regarding two-coil experiments was used to populate a modified version of the TETRAD system. As an example of how this works, eight connections derived from two-coil experiments were used:

These connections are all into lM1, so there aren ‘t any directed edges (connections) that can be chained together, but this allows a simple demonstration of T-Connection applied to this kind of data, since we can *condition* on various sets of variables. There are 16 unconditional T-Connections, each ordered pair of lM1 with the other 8 regions. These are just the edges in the data, all into lM1. However, we can condition on lM1, the common child, on pairs of other regions; if we do this, we get many T-connecting paths, including: This shows what ‘s expected, that all pairs of parents of lM1 are conditionally associated–that is, if you were to experimentally control activity in lM1, you would find the parents of lM1 being associated. The single-edge paths are also included here, interestingly, which is correct even if non-intuitive.

## 4. Discussion

### 4.1. Recap

TMS studies that directly or indirectly address interregional brain connectivity are common, and becoming more so, with the advent of TMS-evoked potentials. There is value in organizing the large amount of data generated into a form that allows the scientific community to explore it with the goal of generating models and mechanistic hypotheses. Machine reading of the literature has thus far been best for extracting *concepts* rather than data; extraction of data required significant expert human interaction. Accordingly, data extracted from papers by humans was entered into a *flat* database that served as input to a graph database (planned to be available at https://musc.edu/ncnm4r/brainstimconn although possibly elsewhere.) The diversity of TMS-connectivity studies requires flexible structures for representation and the storage of data on nodes allows easy expansion if additional characteristics related to a connection need to be added.

### 4.2. Proposal for a database of published TMS connectivity studies

We propose scaling up the database to include all publications that provide insight into interregional brain connectivity. This would mean incorporating the most straightforward studies of conditioning M1 output (MEP) with stimulation of another region as well as more complex studies that relate behavioral effects of TMS to interregional brain networks. Rather than putting all the data into one type of database it should be possible to have overlapping databases that all reference the same brain areas, even if the way connections are described differ.

#### 4.2.1. Two-level database: Data and Models

In thinking about this work, we have realized that there are two levels of network data – basic data about a connection or effect, and models of network structure and causal relationships. These different levels exist in the same space of interregional connectivity, and ideally would be the same, with data supporting every connection and its functional role. However, modeling approaches will need to reflect conflicting and uncertain data, so the model and data graphs will not perfectly overlap. This can be handled by keeping the two types of databases separate, with model databases derived from the representation of extracted data.

### 4.3. Comparison to other data aggregation approaches

in genetics, and how they helped scientific advancement The foremost aggregation of biological data is in the area of genomics, where large public domain databases store vast amounts – on the order of 10^12^ sequences – of genome information [30]. This is not surprising, given the digital-like structure of nucleotide sequences. But while the tools for search and comparison are also highly developed in this area, they do not represent networks, even if modeling approaches are used. In contrast, there has been slower growth in the field of biological networks based on manual or semi-automated extraction of relationships among factors [31].

#### 4.3.1. Potential integration into other databases

We recognize that this effort is not occurring in a vacuum, and there are other databases relevant to brain connectivity. There is also an increasing impetus for sharing raw data and code, including those related to TMS. There is now an automated system for offline analysis of TMS data that can produce uniform output [32] and an experiment control system [33] for a wide variety of electrophysiological experiments,including those using TMS.

### 4.4. Remaining Challenges

Besides the challenges that we have already encountered, expansion of the database project will undoubtedly bring new ones. There are issues related to ownership of information and authentication/privacy of users. Not all publications are accessible to all readers, so issues of fair use of extracted information may arise. While data sharing and openness in general about research are trending upwards, there may be some scientists unhappy about either use of data, or knowledge by others of data retrieval. These issue may become elevated with incorporation of raw data. Nevertheless, as mentioned above, there are numerous examples of similar projects in related brain connectivity domains, so solutions to address these issues exist.

#### 4.4.1. Causality Concerns

Questions of causality are fundamental to science but discussions about causal models are relatively rare. Nevertheless there is a literature addressing thinking about causality in epidemiology, particularly. and biomedicine in general e.g. [34]. Besides the work cited above, evaluation of timing in highly and recurrently connected neural elements will depend on more than just including timing from TMS experiments. Even if TMS effects are ascribed to the most direct connection being considered, there are almost always mutliple pathways between any two nodes in the CNS. So we don ‘t want to overstate the value of TMS data, while we see the value of stimulation methods in providing information that informs causal models in a dynamic way.

## 5. Acknowledgements

Thanks to Kara Nicole-Simms Bocan, PhD for initial database coding, and students, Evan W. Becker, Breanna Donaghy, Sahara Grinkewitz, Tina Marino, and Alexandra Anton Mahfoud, for data extraction and feedback.

## 7. CRediT authorship contribution statement

**George Wittenberg:** Conceptualization, Methodology, Investigation, Formal analysis, Visualization, Writing – original draft, Writing – review & editing, Supervision, Funding acquisition. **Xiaoqi Fang**: Conceptualization, Methodology, Visualization, Writing – review & editing, Software. **Bryan Lee**: Investigation, Data Curation. Writing - Review & Editing. **Nataša Miškov-Živanov**: Conceptualization, Methodology, Writing – original draft, Writing –review & editing, Supervision. **Harry Hochheiser**: Conceptualization, Methodology, Writing - Review & Editing. **Layla Banihashemi**: Writing - Review & Editing. **Michael Vesia**: Writing – review & editing. **Joseph Ramsey**: Conceptualization, Methodology, Visualization, Writing – review & editing, Software, Formal analysis.

